# Identification of motor unit discharges from ultrasound images: Analysis of in silico and in vivo experiments

**DOI:** 10.1101/2024.01.18.576300

**Authors:** Robin Rohlén, Emma Lubel, Dario Farina

**Author notes:** Correspondence to Dario Farina.

## Abstract

**Objective:** Ultrasound (US) images during a muscle contraction can be decoded into individual motor unit (MU) activity, i.e., trains of neural discharges from the spinal cord. However, current decoding algorithms assume a stationary mixing matrix, i.e. equal mechanical twitches at each discharge. This study aimed to investigate the accuracy of these approaches in non-ideal conditions when the mechanical twitches in response to neural discharges vary over time and are partially fused in tetanic contractions.

**Methods:** We performed an in silico experiment to study the decomposition accuracy for changes in simulation parameters, including the twitch waveforms, spatial territories, and motoneuron-driven activity. Then, we explored the consistency of the in silico findings with an in vivo experiment on the tibialis anterior muscle at varying contraction forces.

**Results:** A large population of MU spike trains across different excitatory drives, and noise levels could be identified. The identified MUs with varying twitch waveforms resulted in varying amplitudes of the estimated sources correlated with the ground truth twitch amplitudes. The identified spike trains had a wide range of firing rates, and the later recruited MUs with larger twitch amplitudes were easier to identify than those with small amplitudes. Finally, the in silico and in vivo results were consistent, and the method could identify MU spike trains in US images at least up to 40% of the maximal voluntary contraction force.

**Conclusion:** The decoding method was accurate irrespective of the varying twitch-like shapes or the degree of twitch fusion, indicating robustness, important for neural interfacing applications.

## Introduction

Human movements are generated by the central nervous system, which provides excitatory inputs to motoneurons in the spinal cord. These inputs are further transmitted from each motoneuron to a group of muscle fibres, forming a motor unit (MU), i.e. a motor neuron and all its innervated fibres. It is possible to record muscle electrical activity from the skin and decode it into the activity of individual motoneurons, i.e., trains of neural discharges. The decoded spinal activity can be used for neural interfacing purposes, such as prosthetic control.

The electrical signal generated by the active muscle fibres (electromyographic signal, EMG) is attenuated by the non-homogeneous tissue between the muscle fibres and the electrodes. This limits non-invasive EMG to only recording from fibres close to the skin and intramuscular EMG to monitoring activity very close to the electrode sites. In both cases, the analysis of muscle electrical activity is relatively local.

In addition to the electrical activity, the displacements of muscle fibres in individual MUs can also be measured. Given the electromechanical coupling, an electric impulse corresponds in the mechanical domain to a shortening and thickening of the muscle fibres (mechanical twitch).

However, the twitches have fundamentally different characteristics to the electrical domain’s action potentials. For example, the duration of a mechanical twitch is much longer than that of an action potential. Indeed, a mechanical twitch is usually longer than the interval between neural discharges, leading to partial fusion of twitch responses (Fig. 1). Typical force traces from the tendon under different firing rates for a slow and fast twitch MU (using simulation) are shown in Fig. 1A and 1B, respectively. The firing rate is positively correlated with the degree of fusion.

**Figure 1.**
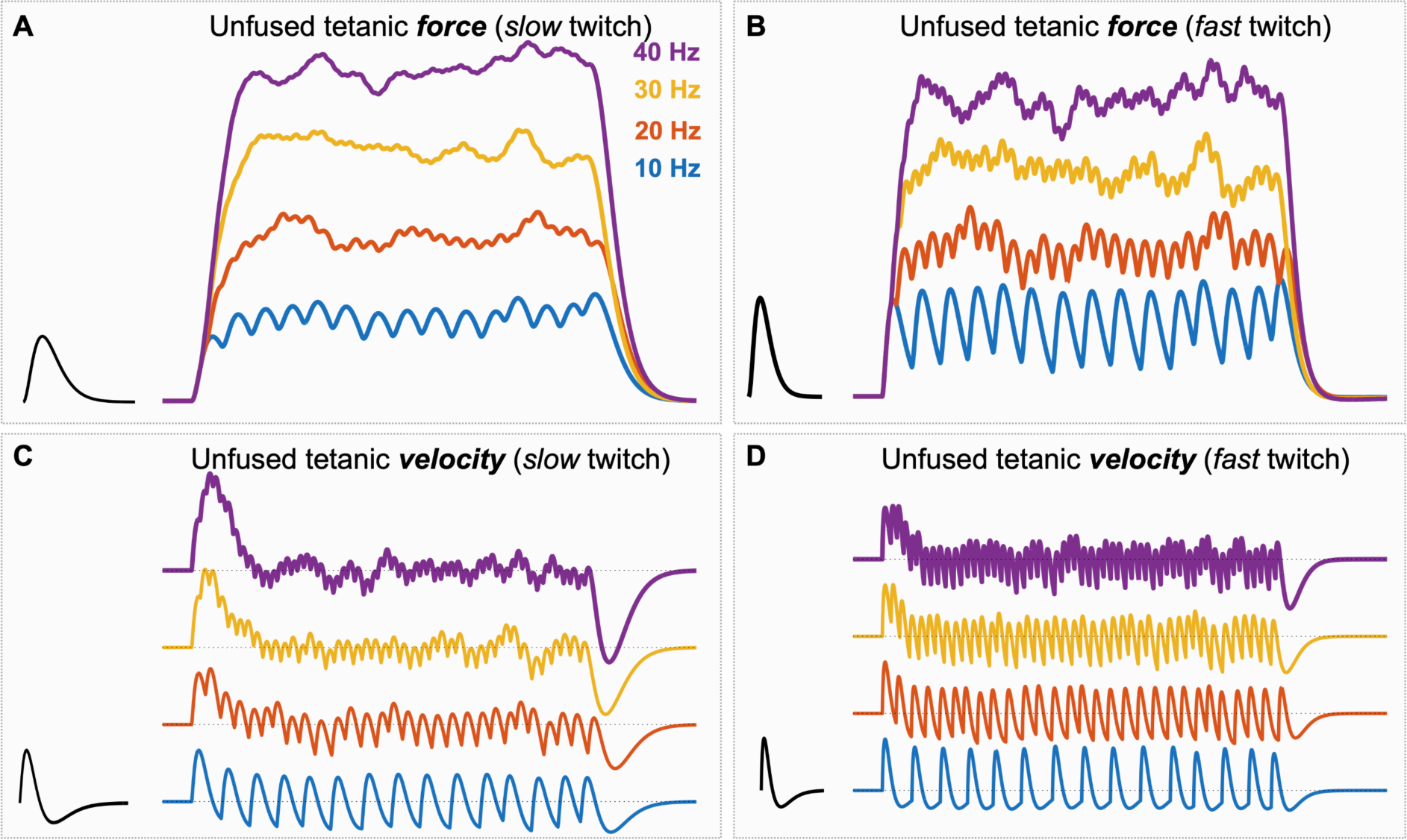
The twitch duration is usually longer than the time between neural discharges leading to (partial) fusion or (unfused) tetani, where the firing rate positively correlates with the degree of fusion. The force traces from the tendon under different firing rates for (**A**) a slow and (**B**) a fast twitch motor unit (MU) differ. (**C**-**D**) Alternatively, the unfused tetani can be described as the velocity of the thickening or the axial movement of the muscle fibres. These plots were produced using simulations with Gaussian-distributed variability in the inter-spike interval (ISI) times, with an ISI coefficient of variation of 10%. The twitch responses were generated using the mathematical equation described in the Methods section, with *V_max_* = 1, *T_max_* = 25, and *T_c_* = 75 in the slow twitch case and *V_max_* = 1, *T_max_* = 10, and *T_c_* = 40 in the fast twitch case.

From an alternative perspective, the unfused tetani can be described as the velocity of the thickening or the axial movement of the muscle fibres (Fig. 1C and 1D). The velocity traces are associated with the ultrasound (US) measure of MU activity. US images can be decoded into individual sources by blind source separation (BSS) methods, either with linear instantaneous models [1,2] or convolutive models [3]. Specifically, convolutive BSS showed high accuracy in identifying discharge series of individual MUs in superficial and deep parts of the tibialis anterior (TA) muscle [3]. However, BSS assumes a stationary mixing matrix, i.e. equal twitches at each discharge for individual MUs. This assumption may not be entirely met [4,5], for example, because of non-linear summations of twitches from other MUs [6]. It is currently unclear how varying twitch waveform shapes influence the accuracy of the current decoding algorithm. Also unclear is whether the degree of fusion of the tetanic contraction influences the accuracy of the decoding algorithm.

This study aimed to investigate the accuracy of a convolutive BSS method [3] in estimating MU discharge times from US images that comprise varying twitch-like shapes in response to neural discharges of each MU, and a varying degree of fusion of the tetanic contraction. For these purposes, we performed an in silico experiment using current knowledge about the experimental spatial distributions [7] and temporal characteristics of MU behaviour [8]. Using the ground truth enabled us to study the decomposition accuracy for changes in simulation parameters, including the twitches, territories, and motoneuron-driven activity. Finally, we explored the consistency of the findings from the in silico experiment with an in vivo experiment on the TA muscle at varying contraction forces.

## Methods

### In silico experiment

To understand how different degrees of tetanic fusion and varying twitch-like shapes within a MU change the spike train accuracy using the convolutive BSS method [3], we simulated image sequences (40×40 mm transverse images over time; 100×100 pixels) corresponding to the axial velocity field in US images during isometric contractions similar to previous simulation approaches [2]. The model of Fuglevand and colleagues inspired us to develop a similar motoneuron-driven model with a recruitment and rate coding organisation [9]. To make the model as realistic as possible, we adapted the functionally-based model to the experimental features of MU twitch profiles [8] and MU territory distributions [7]. We also added three noise levels: no added noise, 20 dB, and 0 dB white noise. In this way, we could quantify the precise change in accuracy after a parameter change associated with the twitch waveform shapes, territories, noise, and discharge rate. Below, we describe each part of the simulation model in detail.

### Motoneuron pool

The simulation model comprised a pool of *n* = 200 motoneurons where the time of the *j*th discharge (spike) of the *i*th motoneuron was modelled as:

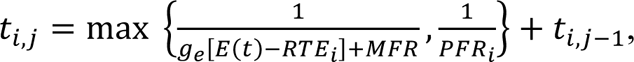

given that *E*(*t*) ≥ *RTE_i_* holds, where *E*(*t*) is an excitatory drive function representing the effective synaptic current, *RTE_i_* is the *i*th recruitment threshold excitation (RTE), i.e., the minimum level of excitatory drive required to initiate repetitive discharges, *MFR* is the minimum firing rate once a motoneuron was recruited, *PFR_i_* is the maximal firing rate of the *i*th motoneuron, and *g_e_* is the gain of the excitatory drive-firing relationship, which was assumed linear.

In this work, we set the gain and the minimum firing rate to fixed values, i.e., *g_e_* = 1, *MFR* = 8 Hz, and the RTE for the *i*th motoneuron as 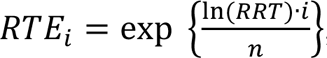, where *RRT* = 30 denote the range of recruitment threshold values, and the peak firing rates *PFR_i_* = *PFR*_1_ − 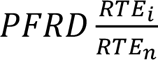 where the peak firing rate for the first motoneuron was set to be *PFR_i_* = 45 Hz. The difference in peak firing rates between the first and last recruited motoneurons was *PFRD_i_* = 10 Hz, as in the original model [9]. Furthermore, we imposed inter-spike interval (ISI) variability onto the model by adding a Gaussian-distributed randomly generated sample with a coefficient of variation equal to 15% for each discharge.

The excitatory drive function *E*(*t*) was set to be a 30-s-long trapezoid function, i.e., a linear increase for the first 5 s from 0% to either 10, 30, 50, 70 or 100% of the maximum excitatory drive, and then a stable level of force for 20 s, and finally a linear decrease down to 0% again. In Fig. 2, we illustrate the firing rate as a function of the excitatory drive (Fig. 2A) and the neural drive to muscles for the first 10 s of a 20% trapezoid function using the above-mentioned parameters (Fig. 2B).

**Figure 2.**
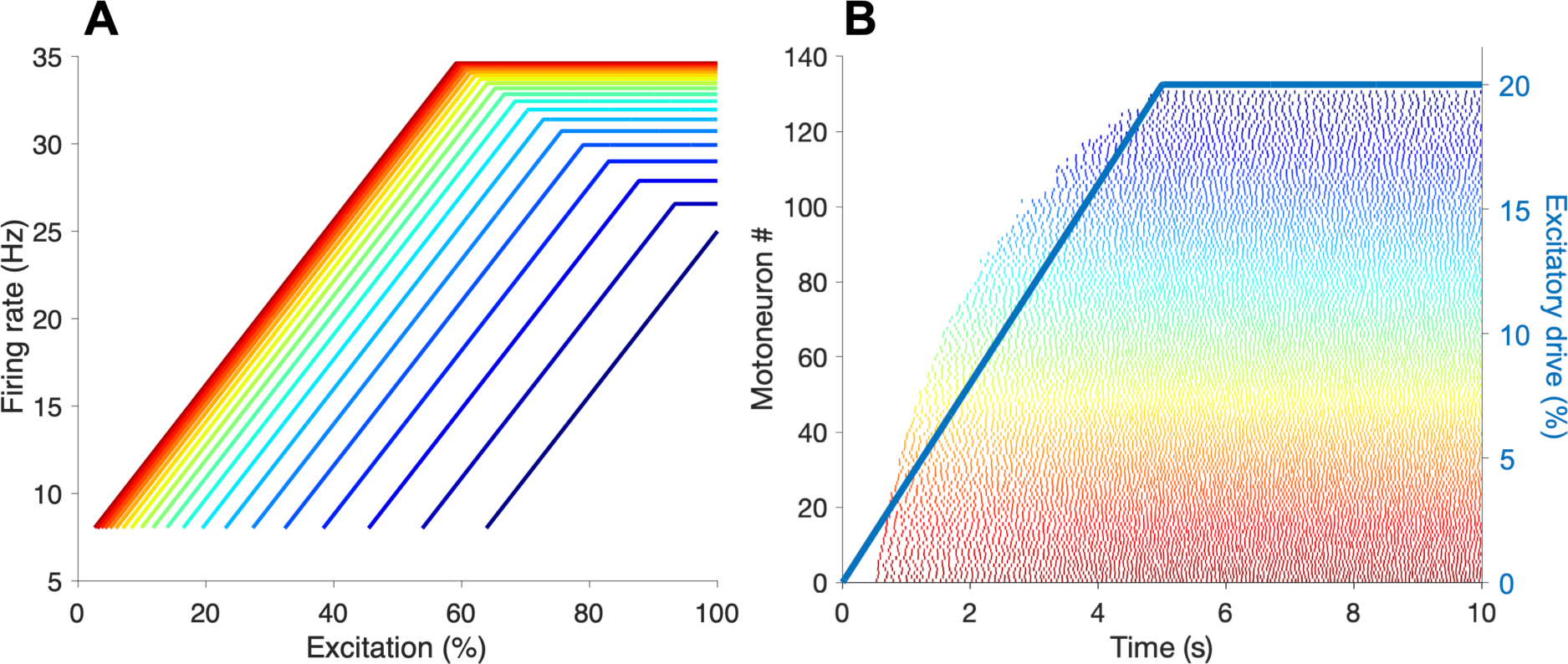
The simulation model modelled the firing rate as a function of excitation max (**A**) with 8 Hz firing rate once any unit is recruited. The maximum firing rate differs between units, and it was 35 Hz for the first and 25 Hz for the last, meaning all units had maximum firing rates between 25 and 35 Hz. (**B**) An example of the spike trains of the recruited motor unit (MU) pool for an excitatory drive curve shaped like a trapezoid with a flat section at 20%.

### MU twitch profiles

We refer to the MU twitch profile as the mechanical response(s) for each discharge of a specific motoneuron. In contrast to typical non-negative twitch profiles in the force or displacement domain, we consider velocity twitch profiles since we intend to simulate image sequences corresponding to the axial *velocity* field during isometric contractions. Here, we used an analytical equation inspired by a previous study where the mechanical force production of electro-stimulated single MUs in the medial gastrocnemius muscles of rats was modelled [4]. Each velocity MU twitch profile can be described as:

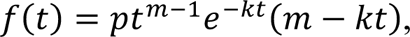

where *t* ≥ 0 and where the MU twitch profile can be described by three parameters: *V_max_*, *T_max_*, and *T_c_* (Fig. 3A). Given these constraints, we can find the analytical forms of *m*, *k*, and *p*:

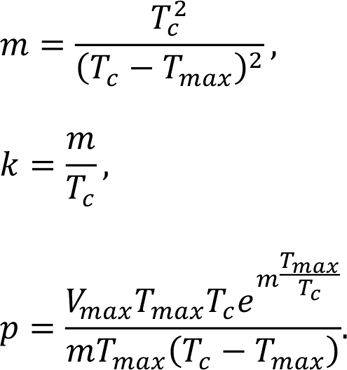

**Figure 3.**
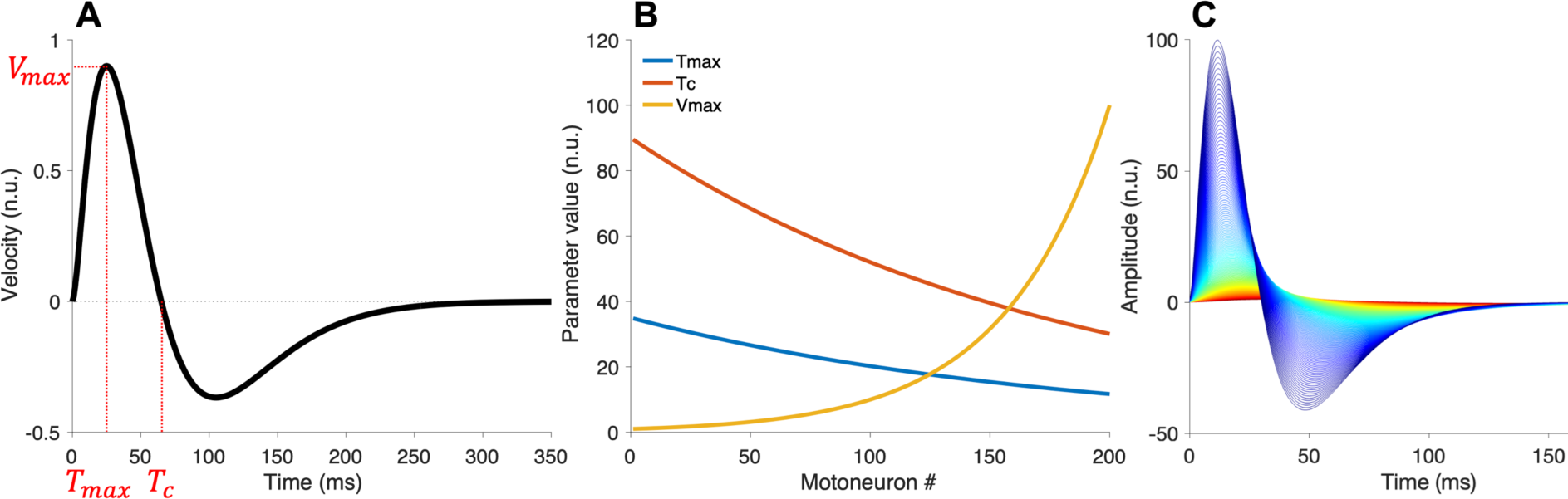
Modelling the motor unit (MU) twitch profiles. (**A**) Each MU twitch profile can be described by three parameters: V_max_, T_max_, and T_c_. (**B**) These three parameters were modelled such that for the first recruited unit the contraction time was the highest and the amplitude (maximal velocity) was the smallest whereas the last recruited was the opposite, i.e., shortest contraction time and largest amplitude. (**C**) All the 200 generated twitch profiles visualised together.

The parameters *V_max_*, *T_max_*, and *T_c_* are only related to the contraction part (the positive values in the velocity domain). The relaxation part (the negative values in the velocity domain) is automatically defined by the value of the same parameters.

The twitch profile parameters were set depending on the order of recruitment to represent the slow and fast units. For the amplitude parameter (*V_max_*), we ensured there was a 100-fold difference in amplitude (*V_max_*) between the first and last recruited MU in the pool and that most units in the pool generated small forces [9]. Therefore, the amplitude for the *i*th MU was set to be 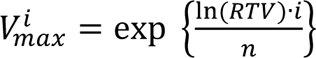 , where *RTV* = 100 denotes the range of twitch velocity values (Fig. 3B). The contraction time (*T_c_*) was set as an inverse power function depending on the amplitude where we considered a 3-fold range in contraction time with the maximal contraction time being 90 ms for the first recruited unit [9]. Thus, the contraction time for the *i*th MU was set to be 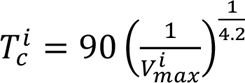 and similarly, the time at maximum velocity for the *i*th MU was set to be 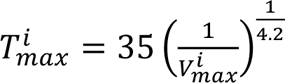 (Fig. 3B). This procedure led to one consistent twitch profile for each MU over the whole time period. In Fig. 3C, we illustrate all the twitches in the MU pool overlayed on each other.

We also generated non-equal twitch profiles within each MU to investigate the decoding algorithm’s accuracy in estimating neural spike trains for varying MU twitch-like shapes in response to neural discharges. The procedure for generating the unequal twitch profile parameters was to draw a random value from a uniform distribution around the equal twitch profile parameters above, i.e., a 50% range around *V_max_^i^* and 20% range around *T_c_^i^* and *T_max_^i^*. Thus, for each twitch profile *j*th discharge (spike) and the *i*th motoneuron, we randomly selected a value from 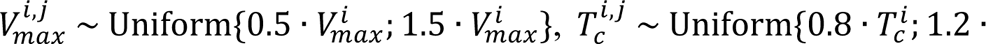 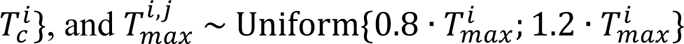.

### MU territories

A MU territory, which comprises the cross-sectional region of the muscle fibres innervated by a motoneuron, was in these simulations a circular region in the 40×40 mm field of view of the muscle cross-section (perpendicular to the fibres). Two parameters described the MU territory. The first parameter was the diameter of the MU territories, which we varied such that the earlier recruited had smaller diameters and the later recruited had larger diameters, i.e., ranging from 3 to 10 mm. Therefore, the territory size for the *i*th MU was set to be *MUI^i^* = 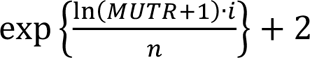, rounded to the nearest integer for practical implementation reasons, where *MUTR* = 10 − 3 = 7 mm denote the range of territory diameter values (Fig. 4A). We defined the MU territories as Gaussian-like distributed velocities similar to experimental observations [7]. In practice, we used weights on the velocities such that the pixels on the centre of the territory were equal to one with a linear decrease in weights as the distance from the centroid increases until zero. The second parameter is the locations of the MU territories. Territories were evenly distributed across the field of view (Fig. 4B), similar to the approach of farthest point sampling [10].

**Figure 4.**
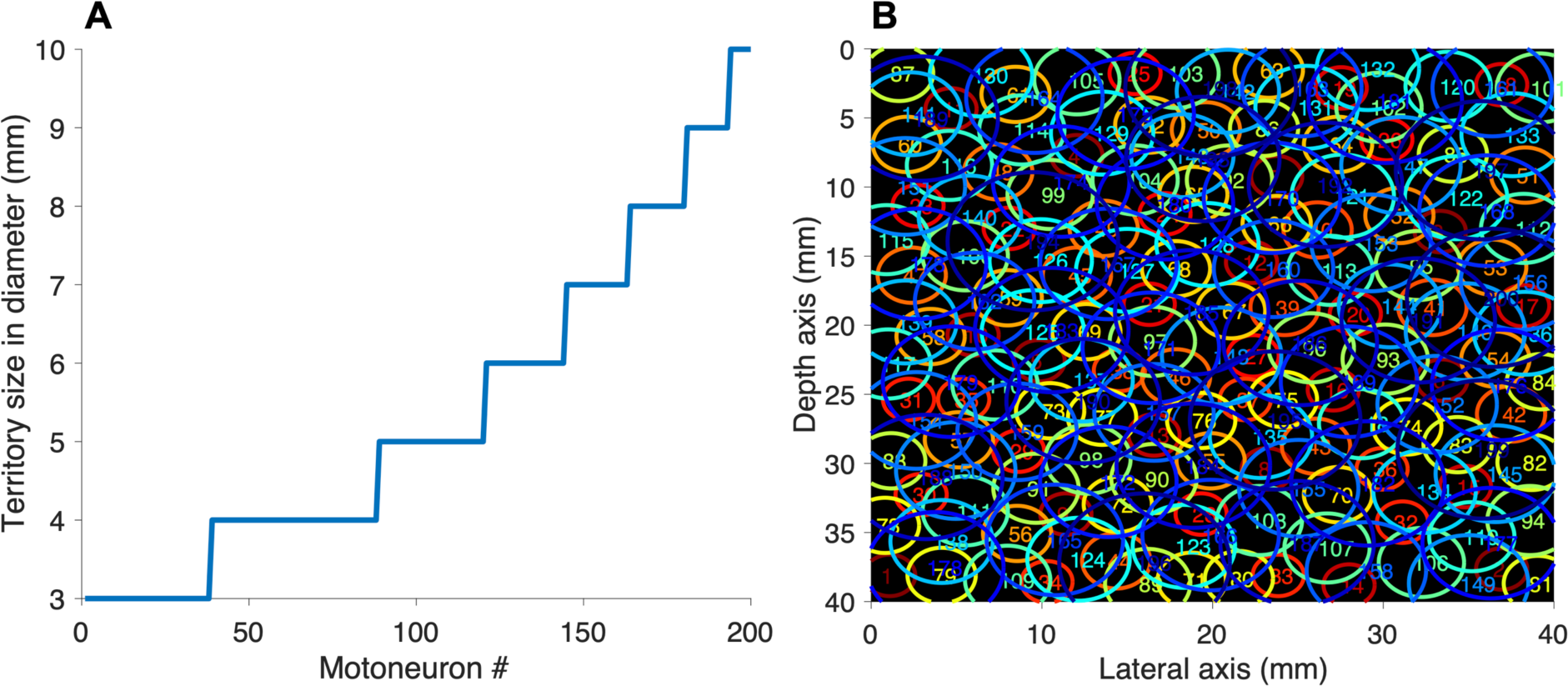
Motor unit (MU) territory modelling. (**A**) The territory sizes in diameter range from 3 to 10 mm, with the earliest recruited having the smallest. (**B**) All the 200 units overlayed on top of each other in recruitment order, from smallest in red to largest in blue.

### In vivo experiment

To explore the consistency of the findings from the in silico experiment, we performed an in vivo experiment comprising a dataset from a healthy 33-year-old man with synchronised ultrafast US, thin-film intramuscular EMG, and force recordings during isometric ankle dorsiflexion contractions of the TA muscle. The study was conducted following the Declaration of Helsinki and was approved by the Imperial College Research Ethics Committee (reference number 19IC5641). Informed consent was obtained from the subject after the study procedures were explained.

After cleaning the skin above the TA muscle, an expert inserted, with the help of a Butterfly ultrasound probe (Butterfly Network, Massachusetts, USA), a thin-film intramuscular EMG array (0.5 mm inter-electrode distance) with 40 platinum electrodes (140×40 µm) [11], a few centimetres below the skin to align it with the image plane of the US probe. Once the needle with the thin-film structure reached the target depth, it was removed, leaving the thin-film with the electrodes inside the muscle. The wires outside the skin were secured with tape, and the leg was secured in an ankle dynamometer (OT Bioelettronica, Torino, Italy).

The participant performed two maximum voluntary isometric (MVC) contractions, the highest one being the MVC reference. Then, the US transducer (L11-4v, 7.24 MHz centre frequency) was placed on the skin with a custom-built probe holder to secure it. Two repeats of five force-level trapezoid ramps were performed at 5%, 10%, 20%, 30%, and 40% MVC, with 60 s of rest between each repeat. Each ramp had a 40-s plateau with 5-s recruitment and de-recruitment ramps. Once the participant reached the plateau, a 30-s ultrafast US recording was performed.

The EMG signals were sampled at 10240 Hz, bandpass filtered at 10-4400 Hz, and A/D converted with 16-bit resolution using a Quattrocento amplifier (OT Bioelettronica, Torino, Italy). The force data were fed through the Forza force amplifier (OT Bioelettronica, Torino, Italy) into the auxiliary port of the Quattrocento. The ultrafast US data were recorded (28.96 MHz sampling frequency) with the Vantage Research Ultrasound System (Verasonics Vantage 256, Kirkland, WA, USA) at a frame rate of 1 kHz using single-angle plane wave imaging. We aligned the data offline after sampling a trigger pulse in the Quattrocento from the Verasonics system via an Arduino Uno [12]. Finally, the recorded EMG signals were decomposed to obtain MU spike trains using an open-source software [13].

### Convolutive blind source separation model of US images

Here, we briefly describe the convolutive BSS model and implementation for decoding US-based velocity images to spike trains based on the original work by [3]. The model represents the axial velocity in the *i*^tA^ pixel as *x_i_*(*t*), *i* = 1, 2, … , *m* over time *t* ∈ [0, *T*] as:

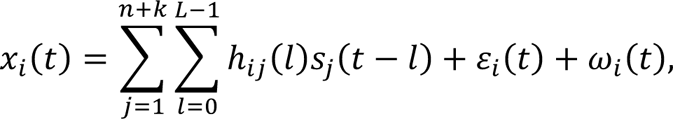

where *n* is the number of MUs, *k* is the number of non-MU sparse sources such as blood vessel pulsations, *s_j_*(*t*) is the *j*^*t*A^ source at time *t*, *h_ij_*(*l*) is the impulse response for the *i*^*t*A^ pixel and *j*^*t*A^ source with (finite) duration *L*, *ε_i_*(*t*) is the additive non-white noise such as bone movement, and ω_i_(*t*) is the additive white noise at the *i*^*t*A^ pixel. As explained in the original work, we assumed *ε_i_*(*t*) to be negligible due to stable isometric contractions [3]. Moreover, the sources are sparse with mostly zeros and a few ones, and the impulse response (twitch profile for a specific source) is the same for each discharge. In this work, we intend to explore how important the latter assumption is for the method’s performance.

### Convolutive blind source separation implementation

We may represent the axial velocities and sources as ***x***(*t*) = [*x*_1_(*x*),*x*_2_(*t*),..., *x_m_*(*x*)] and ***s***(*t*) = [*s*_1_(*t*), *s*_2_(*c*), ..., *_n;k_*(*t*)], where *m* is the number of pixels and *n* + *k* is the number of MU and non-MU sources. Then, we can reformulate the convolutive mixture to a linear instantaneous mixture of a new set of sources, which contains the original sources and their delayed versions:

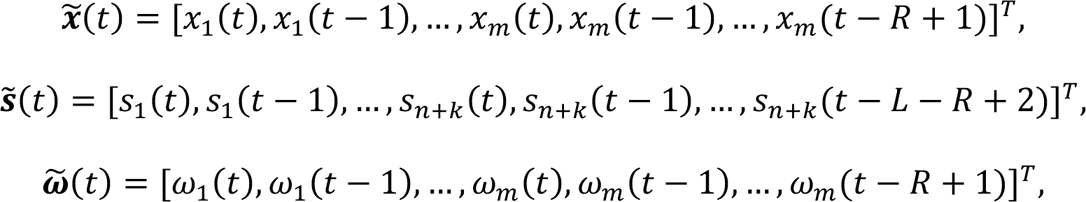

where *L* denote the finite duration of the impulse responses and *R* denote delayed observations as described in the literature [3,14]. In matrix form, we have:

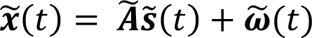

This representation enables a simple implementation using the fixed-point iterations approach [3,15] to estimate the sparse sources ***s***(*t*) by finding the projection (separation) vectors.

For practical reasons, we consider a sliding region of interest (ROI) instead of using a full image sequence as input to the convolutive BSS method, similar to previous works [1,3,16]. Here, we chose non-overlapped ROIs of size 20×20 pixels corresponding to 6×6 mm for the in silico experiments (100×100 pixel images) and slight overlap (for ROIs furthest to the right and bottom) in the in vivo experiments (119×128 pixel images). Next, the ROIs were vectorised, z-scored, and extended with *R* = 20 delayed versions of each observation. Then, the mean was subtracted along each dimension before whitening the extended observation matrix *x*(*t*) using the eigenvalue decomposition of the covariance matrix to induce uncorrelation for time lag zero and unit variance.

After removing stationary, temporal, and spatial white noise by applying a 70% threshold and assigning those eigenvalues smaller than the threshold to noise [3,17], we kept the 100 first eigenvalues for the final stage of the pipeline (except for the no-noise simulation case, where we kept the 400 eigenvalues). As discussed in previous work, the number of eigenvalues used for selection should be subject-specific for optimal performance. However, in this study, using the 100 first eigenvalues provided results close to optimal in both simulations and experimental data. After this step, we applied a fixed-point algorithm with a Gram-Schmidt orthogonalisation [15] to estimate the projection vectors. This estimation was based on maximising the sparsity of the source vector using the same gradient functions as in the original work [3]. Since the estimated sources are not ideal spike trains with zeros and ones, we applied a blind deconvolution method [18] on each estimated source to identify the time instants of the spikes. For more details about the implementation including the pseudo algorithm, see the original work proposing the method [3].

### Quantifying decomposition accuracy

To quantify the accuracy of the convolutive BSS method in estimating neural spike trains upon a varying degree of fusion of the tetanic contraction and varying twitch-like shapes, we calculated the rate of agreement (RoA) between estimated and reference spike trains (ground truth or EMG). The metric is defined as 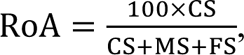, where CS denote the number of correctly identified spikes, MS denote the number of missed spikes, and FS the number of false spikes, given a tolerance window being set to [0,30] ms [18]. We also quantified the time between a correctly estimated spike (CS) and the simulated spike (defined as spike delta) and its variability (spike delta variability).

We defined two separate criteria for a MU match. The first criterion was RoA ≥ 90%. The second criterion was the ratio between the true positives (CS) and the total firings in the ground truth spike train (*n*), i.e., 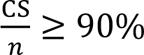 The second criterion is included because it only considers accuracy of identification of ground truth spikes, in contrast to RoA, which is a combined metric including false positives and false negatives.

For the in silico experiments, we selected one estimated source (candidate MU) for each ground truth MU through maximum spike train agreement (criteria above). For the in vivo experiment, we repeated the same procedure but used the spike trains from intramuscular EMG as a reference. However, the uptake area of the 40 electrodes in the thin-film could not access all the units within the field of view of the US transducer (40×40 mm). The criteria to select estimated sources which were not detected by EMG were firing rates above 5 Hz and below 35 Hz, and the coefficient of variation of the ISI to be below 50%. These broad criteria lead to putative false positives, which we removed manually if the sources looked like a mixture of units based on their instantaneous firing rate curves. Finally, to remove duplicates, we calculated the correlation between estimated sources. If the correlation was greater than 50%, we kept the estimated source with the lowest discharge variability while discarding the others.

### Statistical analysis

Descriptive statistics of the rate of spike train agreement, number of MU matches, and temporal correlations were quantified. We tested whether the variation of sources’ amplitudes differed between those from simulated data with equal and unequal successive twitch-like shapes. In addition, we tested whether the variation of sources’ amplitudes differed in experimental and simulated MUs (both for equal and unequal twitches). We set a *p*-value of 0.05 for statistical significance.

## Results

### In silico experiments

We found that a large population of identified MUs had a spike train agreement of at least 90% with the simulated spike trains (Fig. 5). The total number of units with such spike train agreement depended on the level of additive white noise, the percentage of excitatory drive max, and whether the twitches were set to be equal or unequal over time within each MU. The performance was significantly superior without added noise, as expected. Yet, it is useful to include the “null” case in understanding how noise affects the decomposition algorithm. For example, the full motoneuron pool and their neural spike trains can, in theory, be accurately identified if no white noise is present (Fig. 5, blue solid lines). The worst performance was for the unequal successive twitch case and 0 dB white noise (Fig. 5, yellow dashed lines), with ∼60 units found corresponding to ∼65% of the active pool at 10% excitatory max and ∼20 units found corresponding to ∼10% of the active pool at 100% excitatory max.

**Figure 5.**
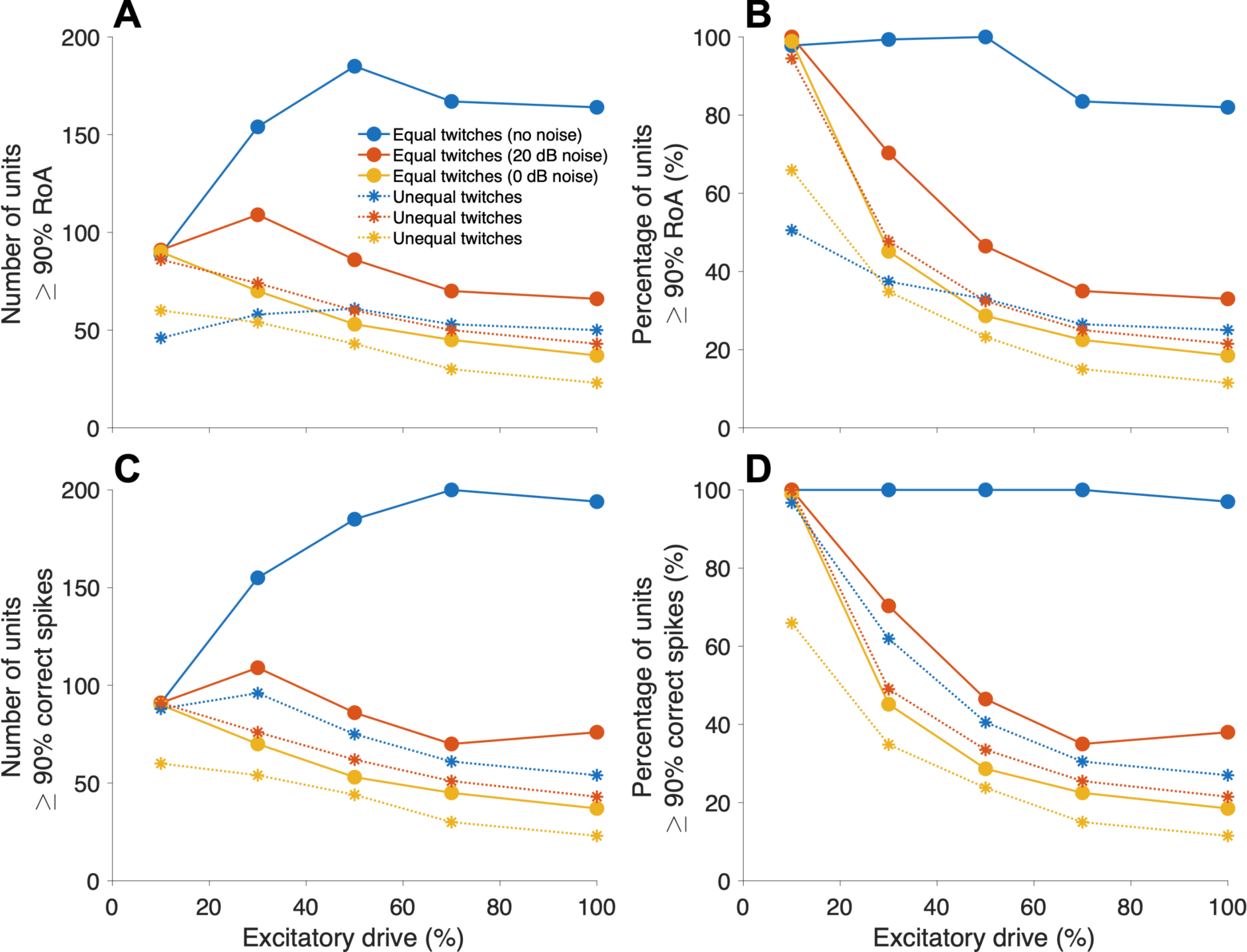
Identification of motor units with accuracy in spike identification of at least 90%. The accuracy is quantified either with (**A**-**B**) the rate of agreement (RoA) metric or (**C**-**D**) percentage of true positives. The absolute values are shown in **A** and **C**, whereas **B** and **D** show the percentages of total number. The number of units with such spike train agreement depended on the level of additive white noise, the percentage of excitatory drive max, and whether the twitches were set to be equal or unequal within each unit.

Although the unequal successive twitch-like shapes affected the number of identified units with high spike train agreement, we found that this was mainly due to the increased number of missed spikes, as illustrated in two cases with 10% (Fig. 6A) and 30% of excitatory max (Fig. 6B) and 0 dB white noise added. The lower amplitudes can explain these missed spikes in the sources for the peak detection method. The lowest possible detectable twitch amplitude depends partly on noise level, but also on the relative amplitudes of the other active units (Fig. 6C). We also found that the latest recruited units in the pool were identified with high spike train agreement (Fig. 6C), showing that the decomposition algorithm easily finds units with the largest amplitude twitches (latest recruited). However, at 10% of excitatory max, all the units could be identified independently from the noise level. Although not simultaneously, we can also study the full MU pool by detecting units at different excitatory max levels (Fig. 6C). Finally, the identified units with spike train agreement above 90% had firing rates ranging from about 8 Hz to 35 Hz (Fig. 6D), suggesting that the method maintains high accuracy even for high degrees of twitch fusion.

**Figure 6.**
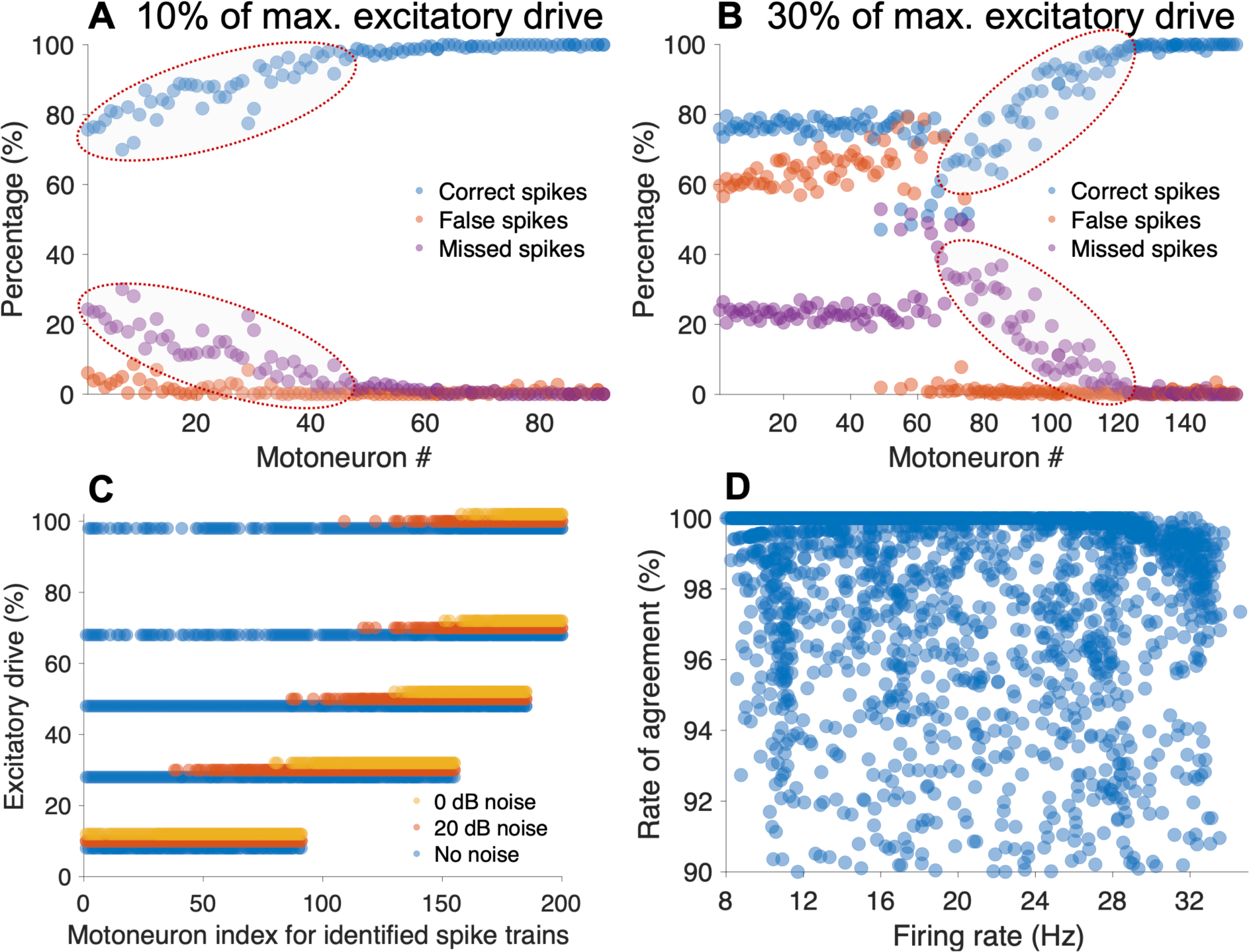
The unequal successive twitch-like shapes reduce the number of identified units with high spike train agreement mainly due to the increased number of missed spikes, as illustrated in two cases with 10% (**A**) and 30% of excitatory max (**B**) and 0 dB white noise added. The latest recruited units in the pool were identified with high spike train agreement (**C**), showing that the decomposition algorithm easily finds units with the largest amplitude twitches. Although not simultaneously, we can also study the full motor unit (MU) pool by detecting units at different excitatory max levels. (**D**) Out of all the identified units with spike train agreement above 90%, we found they varied in firing rates from about 8 Hz to almost 35 Hz.

Fig. 7 shows an example of a source output (MU #100 at 30% of the maximum excitatory drive) for the equal (Fig. 7A) and unequal successive twitch-like shapes (Fig. 7B) within each MU. We found that when the twitches were equal, the variation of the sources’ amplitudes was small compared to the unequal twitches (*p* < 0.001), which was consistent for most MUs (Fig. 7C). Interestingly, we found that the ground truth twitch amplitudes for MU #150 with unequal twitch-like shapes were highly correlated with the amplitude of the sources (Fig. 7D). This high correlation was consistent for most MUs (Fig. 7E). We found this correlation was also true for other cases, such as when multiplying a common mode signal with the compound signal (velocity images), i.e., the weights and the decomposed sources were correlated (not shown).

**Figure 7.**
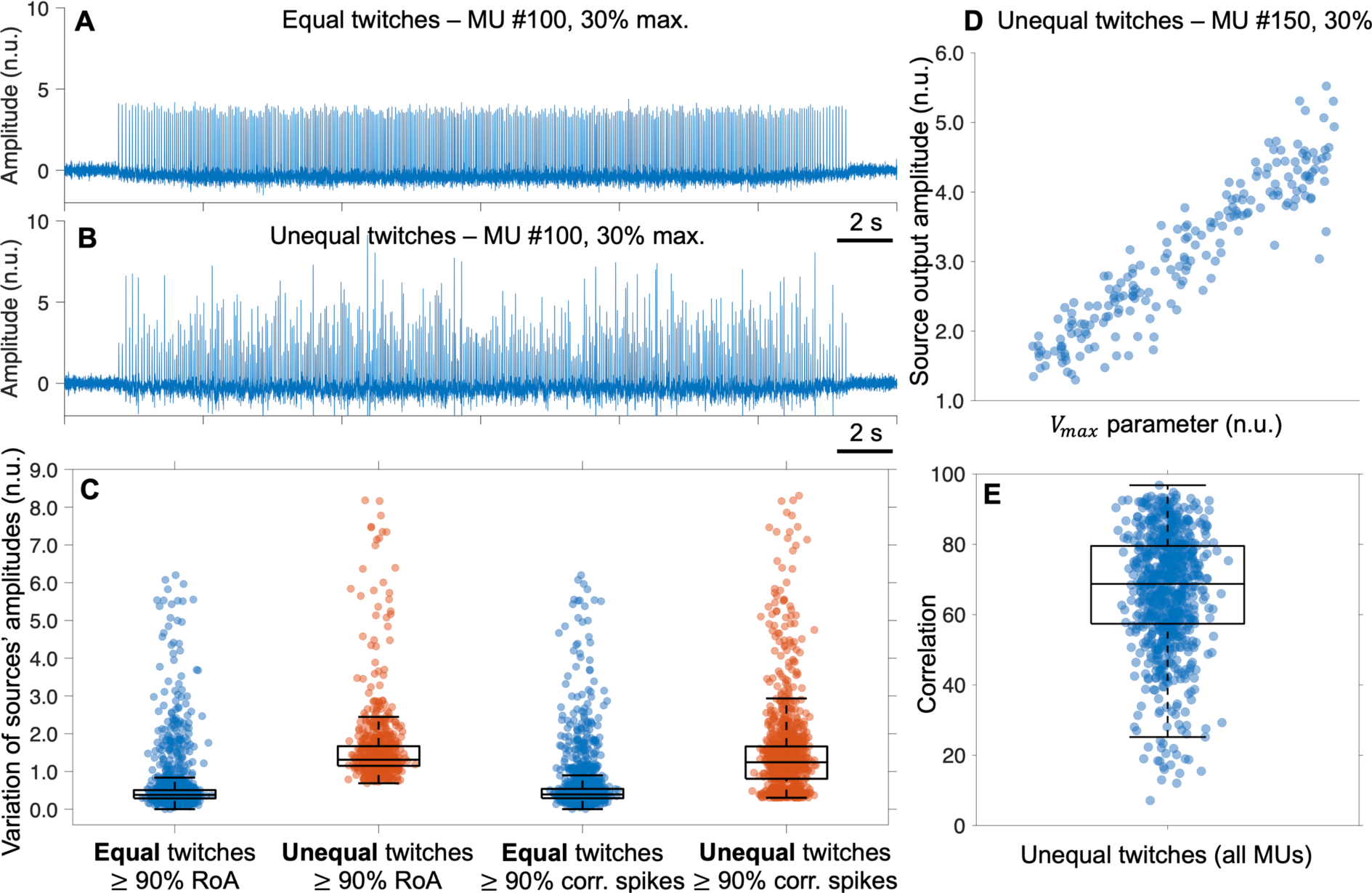
An example of a source output for the motor unit (MU) #100 at 30% of excitatory max for the equal (**A**) and unequal successive twitch-like shapes (**B**) within each MU. (**C**) The variation of the sources’ amplitudes for equal twitches is small compared to the unequal ones (p < 0.001). (**D**) An example of a source output for MU #150 with unequal twitch-like shapes showing a high correlation between ground truth amplitudes and the amplitude of the sources (**D**), which was consistent for most MUs (**E**).

Given the identification of many MUs with a high spike train agreement, the delay between the ground truth and estimated spikes (spike delta) was, on average, ∼12 ms, ranging from about 1 to 25 ms depending on the MU, with no difference between equal or unequal twitches (Fig. 8A) (*p* < 0.001). However, within each MU, the variation in delay across successive twitches was smaller for the equal twitch cases than the unequal twitch cases (*p* < 0.001). The average variation for equal twitch cases was less than 1 ms (Fig. 8B). The variation for unequal twitches was about twice as long (about 2 ms), which was still a limited value.

**Figure 8.**
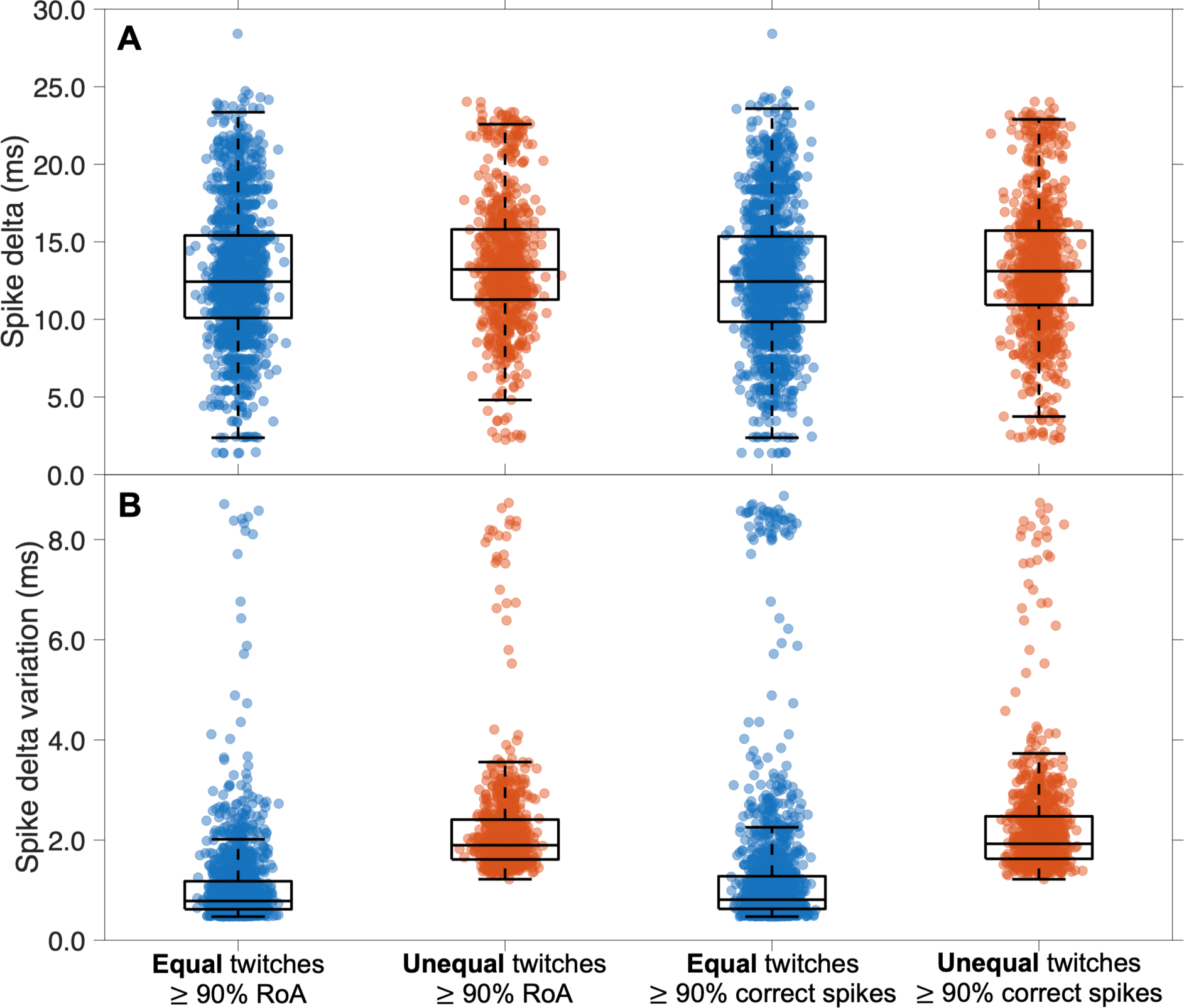
The delay between the ground truth and estimated spikes for the equal and unequal twitch cases did not differ (p < 0.001) and was about 12 ms, ranging from 1 to 25 ms from unit to unit (**A**). Within each motor unit (MU), the variation was smaller for the equal twitch cases than the unequal twitch cases (p < 0.001) (**B**). However, in absolute terms, it is still a low variation compared to the average delay.

### In vivo experiments

To explore the validity of the findings from the in silico experiment, we performed an in vivo experiment on the TA muscle with thin-film intramuscular EMG as a reference. We found good matches between the identified and reference units having a spike train agreement above 90%. Fig. 9A-B illustrates two different matched MUs in the 30-s recordings at 5% MVC. However, after post-processing the data offline, we found that the matched units were on the far left of the US transducer’s field of view, thus the thin-film and probe were not well aligned and the number of matched units between EMG and US was relatively small.

**Figure 9.**
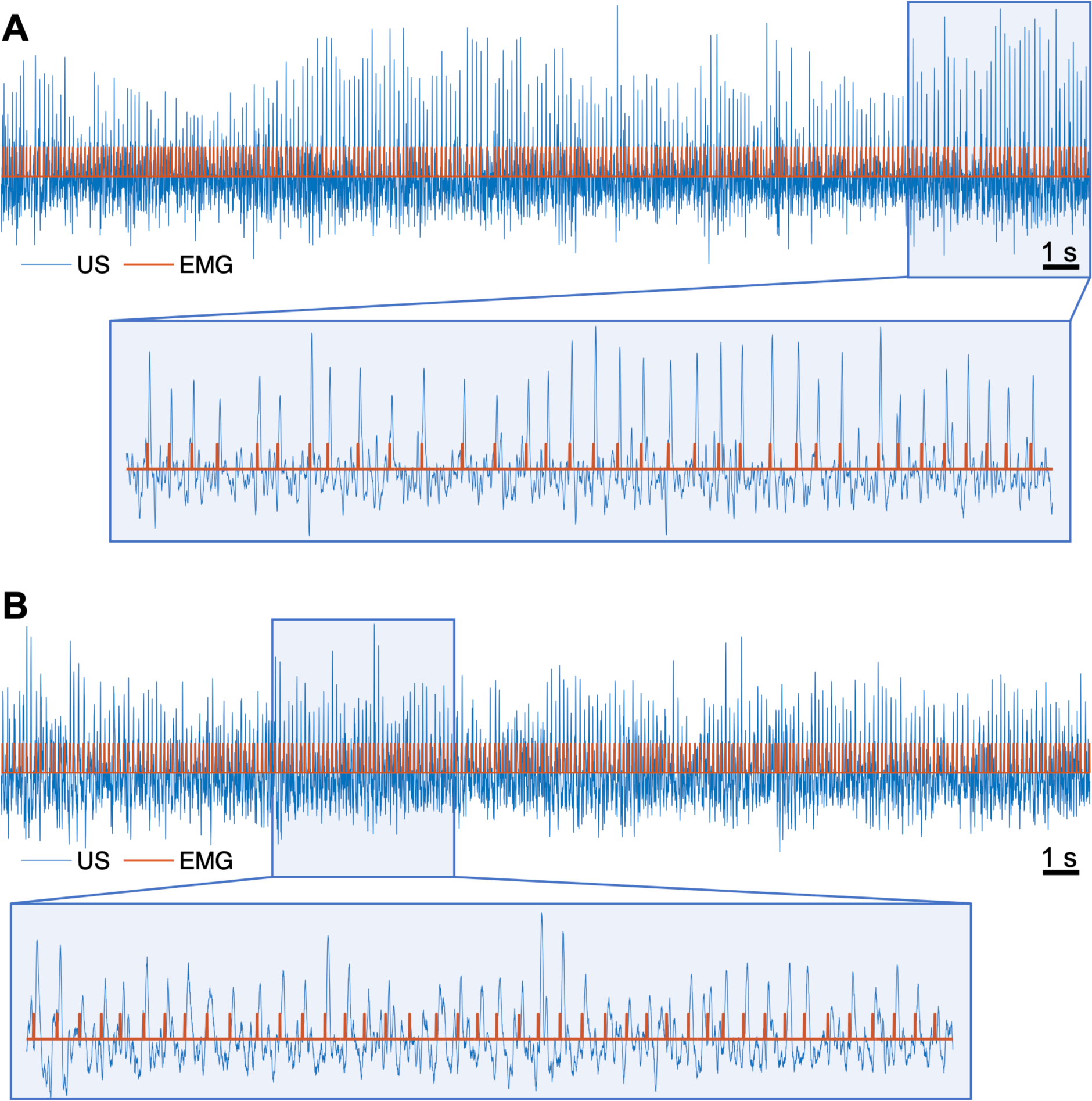
We found good matches between the identified and electromyography (EMG) reference units having a spike train agreement above 90% for two (**A**-**B**) different motor units (MUs) at 5% of the maximum voluntary isometric contraction (MVC).

We found a large number of sources not matched with spike trains from EMG (Fig. 10), being highly similar in “spikiness” to matched units in both simulated and experimental recordings in this study and in previous work [3]. Moreover, these units had firing rates in the range 10–20 Hz, with low variation. Interestingly, we could identify units at all the recorded force levels, including 20% (Fig. 10A-B), 30% (Fig. 10C-D), and 40% MVC (Fig. 10E-F). Note that these sources have not been edited, contrary to what commonly done in EMG analysis [13,19], to avoid any subjective element in the results shown.

**Figure 10.**
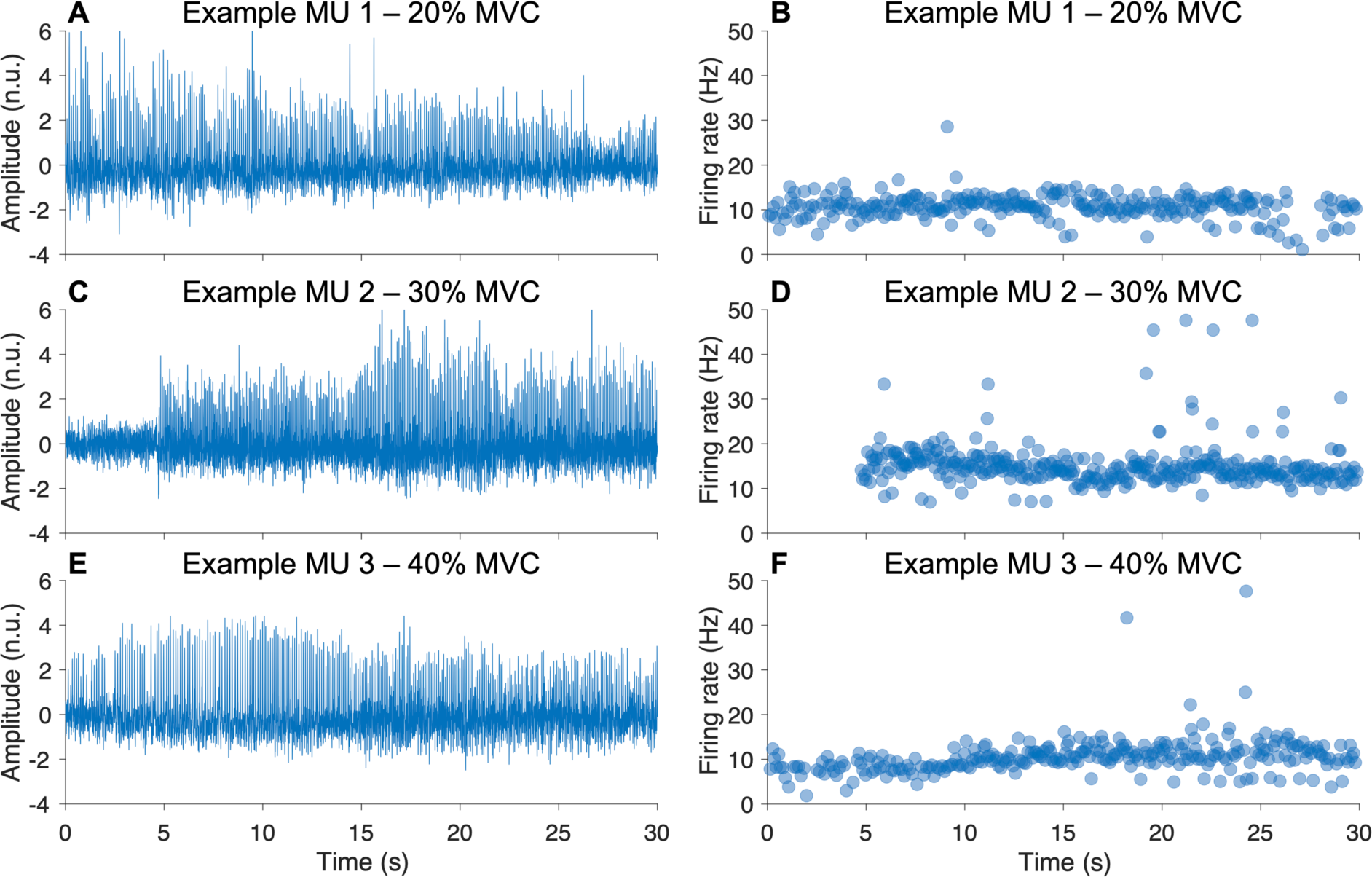
Three examples of sources who did not match with spike trains from electromyography (EMG) but were highly similar in characteristics to matched units in both simulated and experimental recordings. We could identify units at all the recorded force levels, including 20% (**A**-**B**), 30% (**C**-**D**), and 40% MVC (**E**-**F**).

As shown in Figs. 9 and 10, the estimated sources’ amplitudes varied. We found that the variation of the sources’ amplitudes from all the identified units in experimental data was higher than those from simulated MUs at 10-30% of excitatory max with equal twitch shapes and lower than those with unequal twitch shapes (Fig. 11, *p* < 0.001).

**Figure 11.**
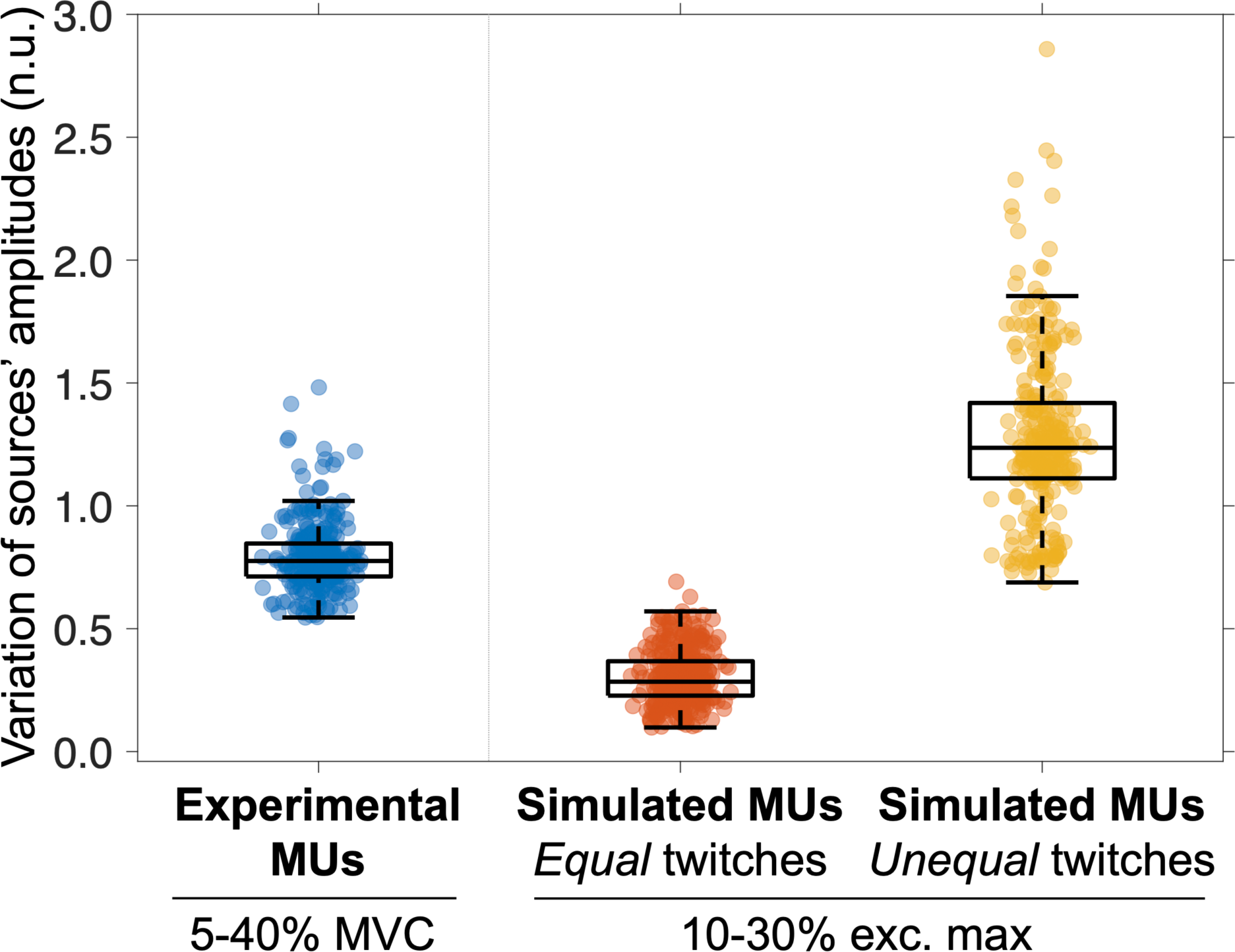
The variation of the sources’ amplitudes from all the identified motor units (MUs) in experimental data was higher than those from simulated MUs at 10-30% of excitatory max with equal twitch shapes and lower than those with unequal twitch shapes.

The identified experimental MUs at 5-40% MVC had varying firing rates, from 8 to 16 Hz, and the distribution centred around 12 Hz (Fig. 12A). In simulations at 10-30% of excitatory max with unequal successive twitch-like shapes, the identified MUs had a similar range of varying firing rates from 8 to 18 Hz with a few up to 20 Hz, and the distribution centred around 11 Hz (Fig. 12B). This indicates consistency between the in silico and in vivo experiments in identifying units with similar degrees of fusion at low-to-medium force and excitability levels.

**Figure 12.**
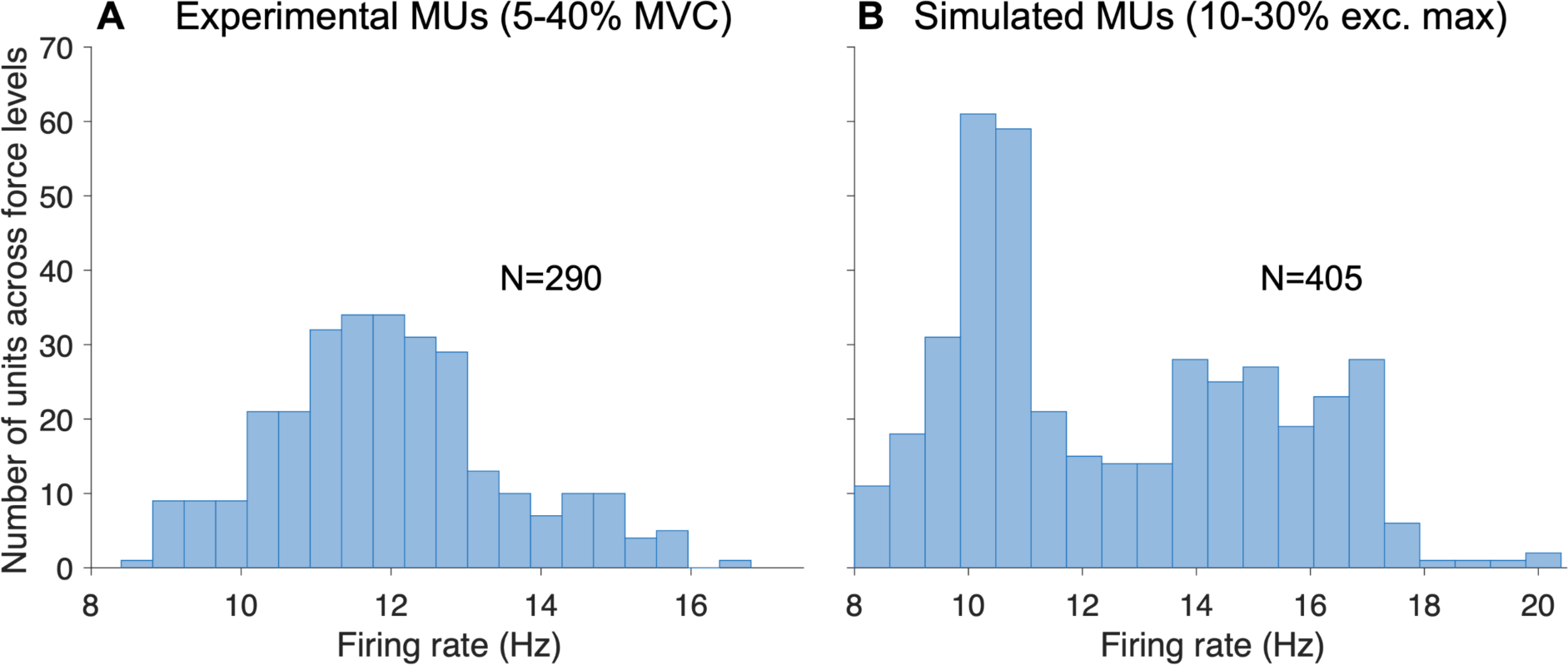
The identified experimental motor unit (MUs) at 5-40% of maximum voluntary isometric contraction (MVC) had varying firing rates, from 8 to 16 Hz, and the distribution centred around 12 Hz (**A**). In simulations at 10-30% of excitatory max with unequal successive twitch-like shapes, the identified MUs had a similar range of varying firing rates from 8 to 18 Hz with a few up to 20 Hz, and the distribution centred around 11 Hz (**B**).

## Discussion

This study aimed to investigate the accuracy of a convolutive BSS method in estimating MU spike trains in US images comprised of varying twitch-like shapes in response to neural discharges of each MU and a varying degree of fusion of a tetanic contraction. We performed in silico experiments and explored the consistency of these findings with an in vivo experiment. We report five main findings. 1) A large population of MU spike trains across different excitatory drives and noise levels could be identified from US, including when the individual MUs had varying twitch shapes over time. 2) The identified MU spike trains with varying twitch waveforms resulted in varying amplitudes of the estimated sources, correlated with the ground truth twitch amplitudes. 3) The identified spike trains had a wide range of firing rates, indicating the method was accurate irrespective of the degree of twitch fusion. 4) The later recruited MUs with larger twitch amplitudes were identified with greater accuracy than those with smaller twitch amplitude. 5) The in silico and in vivo results were highly consistent, and the method could identify MU spike trains in US-based velocity images at least up to 40% MVC.

A large population of MU spike trains across different excitatory drives and noise levels could be identified in silico, including when the individual MUs had varying twitches. The identified MU spike trains with varying twitch-like shapes resulted in varying amplitudes of the estimated sources, which were correlated with the ground truth twitch amplitudes. The twitch amplitude variations impacted the number of units with high spike train agreement (Fig. 5), mainly due to missed spikes (Fig. 6). An adaptive blind deconvolution step could therefore be useful, as discussed previously [3]. However, performing so-called MU editing [13,19] by updating the projection vector will also be required to refine these source outputs. To overcome the post-hoc manual editing for online interfacing purposes, one should consider an automated error correction method using deep metric learning, as recently proposed [20]. Another issue is that current metrics, such as the pulse-to-noise (PNR) ratio [21] and silhouette (SIL) value [14], used for assessing the quality of the decomposition output are not optimal for varying amplitudes of estimated sources, and a new metric, robust to this variation, is therefore needed. However, we found that the amplitude variation of the sources from the in vivo experiments was lower than the simulated variation, thus the simulation results with twitch amplitude variations provide conservative estimates of accuracy.

The identified spike trains had a wide range of firing rates, indicating the method was still accurate at high degrees of fusion. The unfused tetanus of a MU will oscillate around zero, regardless of the firing rate and twitch shape, because the positive and negative parts of a velocity contraction curve approximately cancel each other [8]. In these isometric contractions, where the firing rates are not too high to fuse fully, they still oscillate enough for detectability. However, in some specific subjects or explosive or dynamic conditions, we should expect complete fusion, and thereby, it would be impossible to detect the spike trains indirectly. Another limitation of the technique is that there might be no twitch in some pathological conditions in response to a neural discharge. Nevertheless, in physiological conditions, we conclude that MU spike trains can be identified for most part of the range of discharge rates.

The MUs with larger twitch amplitudes were easier to identify than those with smaller twitches. This finding suggests the method to be biased towards later recruited units, confirming the detectability assumption that a larger MU would have a greater effect on the velocity field of the muscle with respect to a smaller MU, as it involves the movement of more muscle fibres [6]. The question remains whether a 100-fold difference in twitch amplitude (velocity) between the first and last recruited unit is realistic, as we applied this value to velocity twitches adopting it from force twitches [9]. Moreover, as shown in Fig. 6C, studying the full MU pool may be possible by detecting units at different excitatory max levels. Although not confirmed in experimental conditions, this indicates the feasibility of studying the characteristics of the pool of slow and fast MUs that could be important in diagnostics for diseases involving the denervation of a specific MU type, such as ALS and the early denervation of fast MUs.

The in silico and in vivo results were consistent although the simulation model was a linear system and we did not include non-MU motions, such as due to connective tissue, bone movement, etc. Yet, the output of the sources in simulation, which have been processed similarly throughout the study, had characteristics similar to those of experimental recordings, suggesting that the simulations still capture important dynamics in experimental data. There are multiple ways of improving the simulation model. For example, instead of simulating the velocity field directly, one may simulate the US field using Field II [22] and then use axial phase-shift correlation methods to obtain the velocity images. Deep learning-based approaches are more appropriate to make a more authentic simulation model including non-MU activity, e.g., using unsupervised domain-to-domain translation [23].

Physiologically, the displacement field during muscle contractions can be considered a non- linear system, especially due to the connective tissue and the push-and-pull on nearby units and tissue [6,23]. However, this does not imply that we cannot detect all spikes from all motoneurons because we only need to detect the twitch response to a neural discharge, which should be one-to-one. On the other hand, the non-linearities, other non-MU sources, and connective tissue likely affect the detectability of the twitch onsets, e.g., through global or local common mode signals. This could be a reason for missed low amplitude spikes, e.g., Fig. 10A-B after 25 s. We briefly tested this idea by multiplying a pre-defined common mode signal by the image sequence where the amplitudes of the decomposed sources were consistent with the common mode signal. Therefore, we anticipate that removing the common mode in experimental data will improve the MU yield, especially at higher forces. For example, one may use spatiotemporal filtering like singular value decomposition directly on the raw US signals to remove a selected set of singular values [24], as used in, e.g., functional US location microscopy to discard global tissue motion and improve the detection of microbubbles [25].

The convolutional BSS method can identify MU spike trains in US-based velocity images up to 40% MVC. Previous studies using ultrafast US for MU identification and analysis have considered low forces of up to 10% [16,26], 20% [6], and 30% [27], and not with direct identification of the spike trains. This is the first study showing MU spike trains directly identified from US at 40% MVC. Moreover, the in silico experiments suggest the feasibility of identifying MU spike trains at even higher force levels. As shown in Figs. 9 and 10, the decomposed outputs have similar characteristics, i.e., spiky source outputs with varying amplitudes, despite different force levels.

In conclusion, we analysed the accuracy of a convolutive BSS method in estimating MU spike trains in US images comprising varying twitch-like shapes in response to neural discharges of MUs and a varying degree of fusion of a tetanic contraction. By performing in silico experiments and exploring the consistency with an in vivo experiment, we found the BSS method was robust in identifying MU spike trains under varying successive twitch-like shapes, degrees of fusion, and force levels.

## Acknowledgements

The work was supported by funding from the European Union’s Horizon 2020 research and innovation programme under Grant Agreement No.899822, SOMA project. R.R. is supported by the Swedish Research Council (2023-06464), the Swedish Brain Foundation (PS2022- 0021), the Swedish Research Council for Sport Science (FO2024-0003), and the Promobilia Foundation (A23161).

